# Asprosin Neutralizing Antibodies as a Treatment for Metabolic Syndrome

**DOI:** 10.1101/2020.09.15.298489

**Authors:** Ila Mishra, Clemens Duerrschmid, Zhiqiang Ku, Wei Xie, Elizabeth Sabath Silva, Jennifer Hoffmann, Wei Xin, Ningyan Zhang, Zhiqiang An, Atul R. Chopra

## Abstract

Recently, we discovered a new glucogenic and centrally-acting orexigenic hormone – asprosin. Asprosin is elevated in metabolic syndrome (MS) patients, and importantly, its genetic loss results in reduced appetite, leanness and robust insulin sensitivity, leading to protection from MS. Here we demonstrate that anti-asprosin monoclonal antibodies (mAbs) are a dual-effect pharmacologic therapy that targets the two key pillars of MS – over-nutrition and the blood glucose burden. Anti-asprosin mAbs from three distinct species lowered appetite and body weight, and improved blood glucose in a dose-dependent and epitope-agnostic fashion in three independent MS mouse models, with an IC_50_ of ∼1.5 mg/kg. In addition, mAb treatment ameliorated MS associated dyslipidemia and hepatic dysfunction. The mAbs displayed half-life of over 3 days in vivo, with equilibrium dissociation-constants in picomolar to low nanomolar range. This evidence paves the way for further development towards an investigational new drug application and subsequent human trials for treatment of MS, a defining physical ailment of our time.

## Introduction

Obesity and its co-morbidities, such as insulin resistance, hypertension, and dyslipidemia, are omnipresent, affecting nearly a quarter of the world population by some estimates^1^. These conditions, which feed the spread of type II diabetes, coronary artery disease, stroke, nonalcoholic steatohepatitis, and other diseases, are commonly clustered under the umbrella term metabolic syndrome (MS) or syndrome X^1^. MS is a consequence of chronic over-nutrition, turning the evolutionary drive to gather energy from the environment into a liability. As a whole, MS currently exists as an untreatable malady despite decades of basic research and drug development^1^.

Through the study of a rare genetic condition in humans, Neonatal Progeroid Syndrome (NPS, also known as Marfanoid-Progeroid-Lipodystrophy syndrome), we recently discovered a fasting-induced, glucogenic and orexigenic hormone that is the C-terminal cleavage product of profibrillin (encoded by FBN1), and named it asprosin. Its two major sites of action are the liver and the brain. At the liver, asprosin causes a glucogenic effect through a cAMP-PKA dependent pathway^2^. It was found recently to promote hepatic glucose release through the binding of OR4M1, an olfactory G-coupled protein receptor (GPCR) in the rhodopsin family3. In addition, asprosin was shown to bind the mouse analog, Olfr734 with high affinity, and elimination of the receptor considerably reduced the glucogenic effects of exogenously administered asprosin^3^. There is also evidence, that asprosin crosses the blood brain barrier and exerts effects on the hypothalamus^4^. In the arcuate nucleus of the hypothalamus, asprosin directly activates orexigenic AgRP neurons and indirectly inhibits anorexigenic POMC neurons, resulting in appetite stimulation. Patients with Neonatal Progeroid syndrome (NPS), a genetic model of deficiency in plasma asprosin, present with low appetite associated with extreme leanness and robust insulin sensitivity^2,4^. NPS mutations in mice (FBN1^NPS/+^) result in phenocopy of the human disorder, and depressed AgRP neuron activity, which can be restored to normal with asprosin replenishment^3^. Importantly, FBN1^NPS/+^ mice are completely immune to diet-induced MS^4^.On the opposite end of the energy-balance spectrum, patients and mice with MS exhibit elevated plasma asprosin^2-9^.

Based on these observations, we hypothesized that pharmacologic inhibition of asprosin is particularly well suited to the treatment of MS, a condition in need of simultaneous reduction in both appetite and the blood glucose burden. Similar to humans, mice with MS display elevations in plasma asprosin^2,3,9^ making them ideal preclinical models for testing this hypothesis.

To test this hypothesis, we generated three independent monoclonal antibodies (mAbs) that recognize unique asprosin epitopes and investigated their preclinical efficacy and tolerability in the treatment of MS. We specifically dissected the suitability of acute and chronic asprosin neutralization in three different mouse models of MS – mice treated with a high fat diet (diet induced obesity, DIO), mice with genetic leptin receptor mutation (*Lepr*^db/db^), and mice treated with a NASH (Non Alcoholic Steato-Hepatitis) diet. For absolute proof-of-concept, we tested the ability of artificially induced plasma asprosin (adenovirus and adeno-associated virus mediated elevation in plasma asprosin) to raise blood glucose, appetite and body weight, even in the presence of normal chow, followed by rescue of those parameters with immunologic neutralization of asprosin. Our results demonstrate a promising treatment of MS with the use of anti-asprosin mAbs as a targeted, bimodal therapeutic strategy.

## Results

### A single dose of an asprosin neutralizing mAb reduces appetite, body weight and blood glucose levels in mice with MS

We previously showed that adolescent boys with MS have elevated circulating asprosin, and that NPS patients have undetectable plasma asprosin^2,4^. Here we find that a cohort of adult men with MS have significantly elevated plasma asprosin, with levels more than 2-fold higher than those of unaffected age- and sex-matched individuals (Supplementary Fig.1d). This result is consistent a multitude of independent studies^4-9^.

To generate a preclinical model of MS, we exposed C57Bl/6 mice to a high-fat diet for a minimum of 12 weeks (DIO mice). A single dose of an asprosin-neutralizing mAb was sufficient to significantly reduce food intake by an average of 1g/day in the treatment group, but not in mice receiving a control, isotype-matched IgG mAb (Fig 1a). This decrease in food intake was associated with a 0.7g average reduction in body weight over 24h (Fig 1b). In fasted mice, a significant blood glucose reduction was evident as early as 2h post mAb treatment. The effect was most pronounced at 4h and 6h after injection (Fig 1c). Given that mice were without access to food, this result demonstrates that the glucose lowering effect is independent of reductions in food intake.

**Figure 1:**
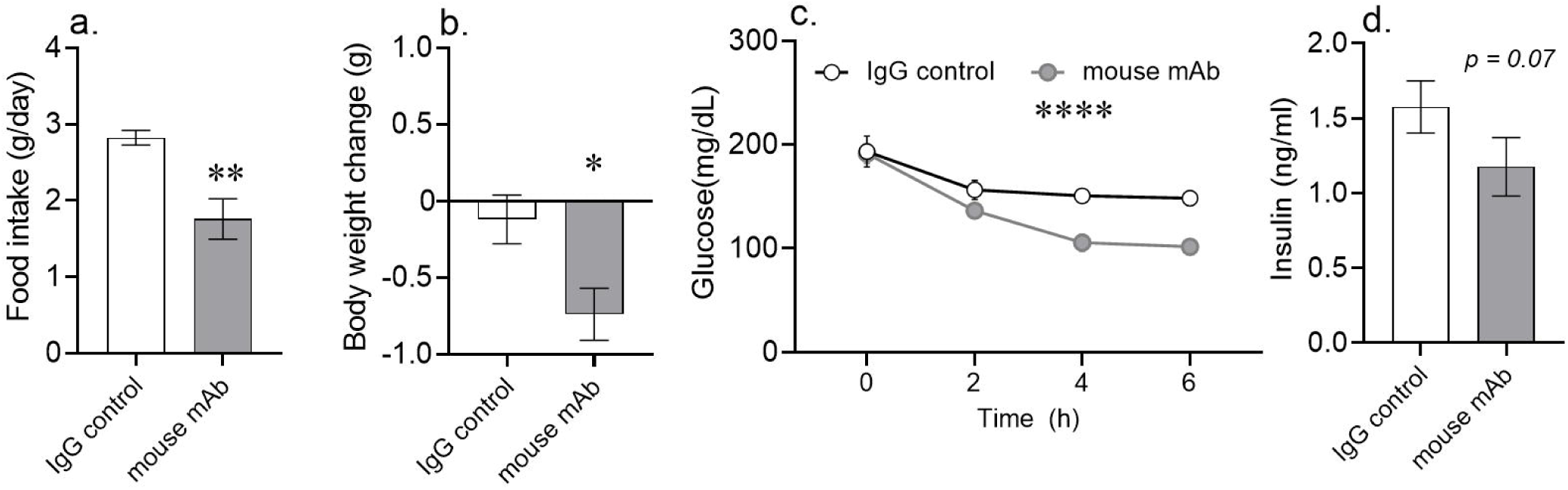
Acute asprosin-neutralization reduces plasma glucose, appetite and body weight in diet induced obese mice. (a-d) Cumulative food intake and body weight change (measured 24h post treatment), baseline blood glucose (measured at hour 2, 4 and 6 post mAb-treatment) and plasma insulin (measured 6h post treatment) were measured after a single dose of anti-asprosin mAb (250µg/mouse) in 16-week-old, male, DIO (diet induced obesity) mice, n = 5 or 6 per group. Note that mice were without food for the duration of the experiment in (c,d), demonstrating that the glucose lowering effect was independent of the hypophagic effect of mAb treatment.

Interestingly, repeating this experiment in lean WT mice (with normal plasma asprosin compared with DIO mice) showed a subtle effect on blood glucose without inducing overt hypoglycemia, and no effect at all on 24h cumulative food intake or body weight (Suppl. Figure 2). Importantly, this result in lean WT mice points towards a clean safety profile of the drug.

**Figure 2:**
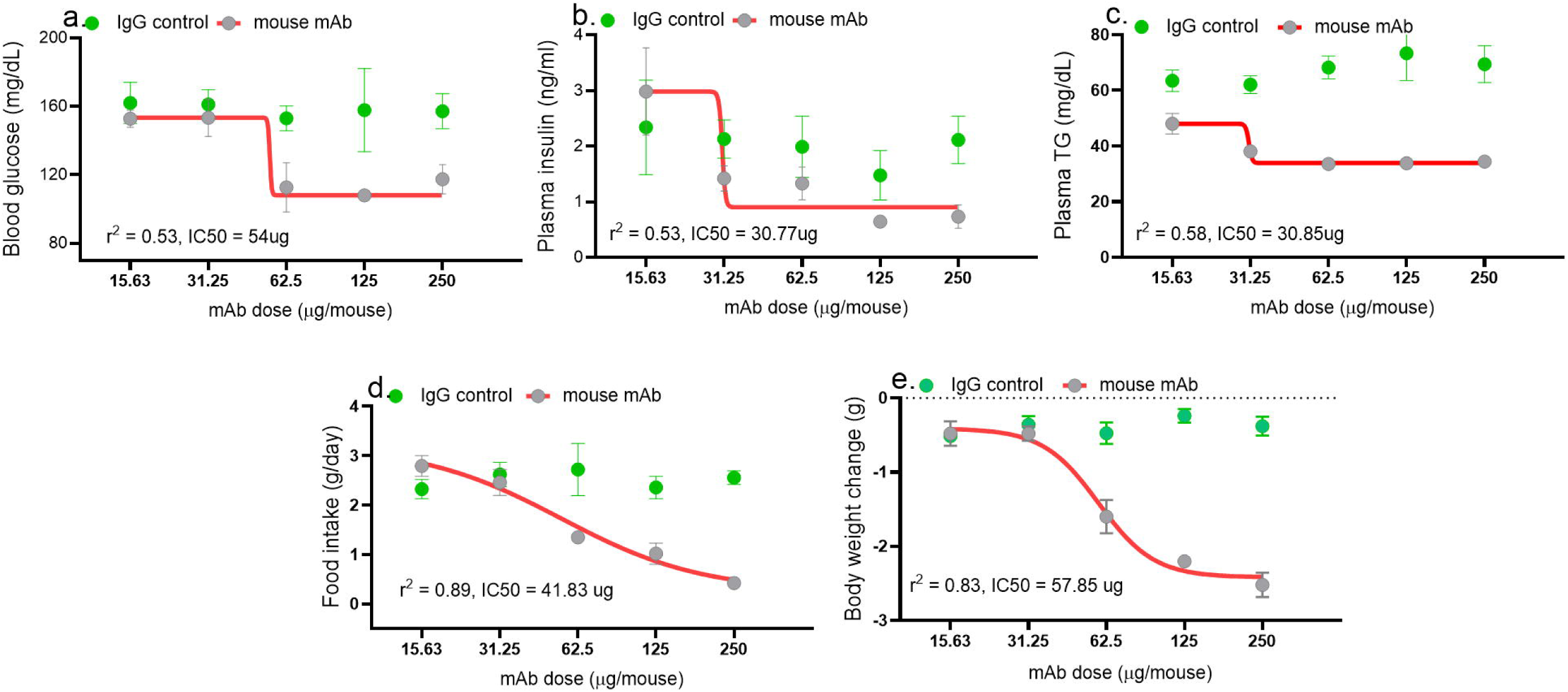
Asprosin neutralization corrects hyperglycemia, hyperphagia and hypertriglyceridemia in a dose dependent manner. Baseline blood glucose (measured 4 h post treatment), plasma insulin and triglyceride (TG; at 24 h post treatment) levels, cumulative food intake, body weight change (measured 24h post treatment), were measured upon injecting increasing dose of anti-asprosin mAb in 16-week-old, male, DIO (diet induced obesity) mice, n = 5 per group. The doses tested were 15.63, 31.25, 62.5, 125 and 250 µg/mouse, corresponding to 0.4, 0.85, 1.64, 3.28 and 6.88mg/kg. Half maximal inhibitor concentration (IC_50_) was determined using a four-parameter non-linear variable slope curve. Asterisk (*) indicate the range of alpha; * p<0.05, ** p<0.01, *** p<0.001, and **** p<0.0001, as determined by student T-test.

### Acute asprosin-neutralization dose dependently mitigates MS

A single dose of the mAb resulted in a significant reduction in blood glucose levels (measured 4h post treatment), plasma insulin and triglyceride levels (measured 24h post treatment), and in 24h cumulative food intake and body weight in a dose-dependent manner in DIO mice (Fig. 2a-e). Half maximal inhibitor concentration (IC_50_) determined using four-parameter non-linear variable slope curve for these three measures was in the range of 30-55 µg/mouse (∼1.5 mg/kg). An isotype-matched, control IgG did not show any effect at all. Interestingly, the dose-response curve shapes were distinct for the five end-points, suggesting that either the potency of asprosin varies among these measures, or they utilize distinct epitopes on asprosin that are differentially affected by this mAb. Further, the plateauing of the mAb effect for body weight, glucose, insulin and triglycerides with higher doses suggests the existence of internal compensatory/buffer mechanisms to protect against drastic reductions in those parameters.

### Viral overexpression of human asprosin induced a hyperphagic, obese, and hyperglycemic phenotype in mice that was rescued by immunologic neutralization

We were interested in the effects of sustained elevation of asprosin in mice fed a normal diet, and in generating tools that obviate the need to employ recombinant proteins of variable activity. To this end, we compared three gain-of-function experiments where we transduced mice with adenoviruses (Ad5) or adeno-associated viruses (AAV8) encoding human *Fbn1* (with native signal peptide) or human cleaved asprosin (with an IL2 signal peptide to promote secretion), and then tested whether viral vector-induced metabolic phenotypes could be rescued with mAb treatment. When compared to mice transduced with Ad5-empty and AAV8-empty control viral vectors, mice transduced with Ad5-*FBN1*, Ad5-*Asprosin* and AAV8*-Asprosin* all exhibited hyperphagia, gained weight and displayed significantly higher baseline glucose and insulin levels (supplementary Fig. 3a-d,f-i,k-n). This metabolic phenotype in all three experiments was coincident with the elevation of human asprosin in mouse plasma in the time it takes to achieve peak expression *in vivo* (supplementary Fig. 3e,j,o)^10,11^. For example, AAV8 vectors typically take up to 50-60 days to achieve peak protein expression *in vivo*^11^. Notably, that is when the mice transduced with AAV8-*Asprosin* first display an increase in body weight (Fig. 3i). In contrast, Ad5 vectors produce a more accelerated peak expression, resulting in quicker increase in body weight (Fig. 3a, 3e). In either case, these three distinct gain-of-function experiments clearly show the effect of asprosin *in vivo* in an experimentally robust and rigorous manner.

**Figure 3:**
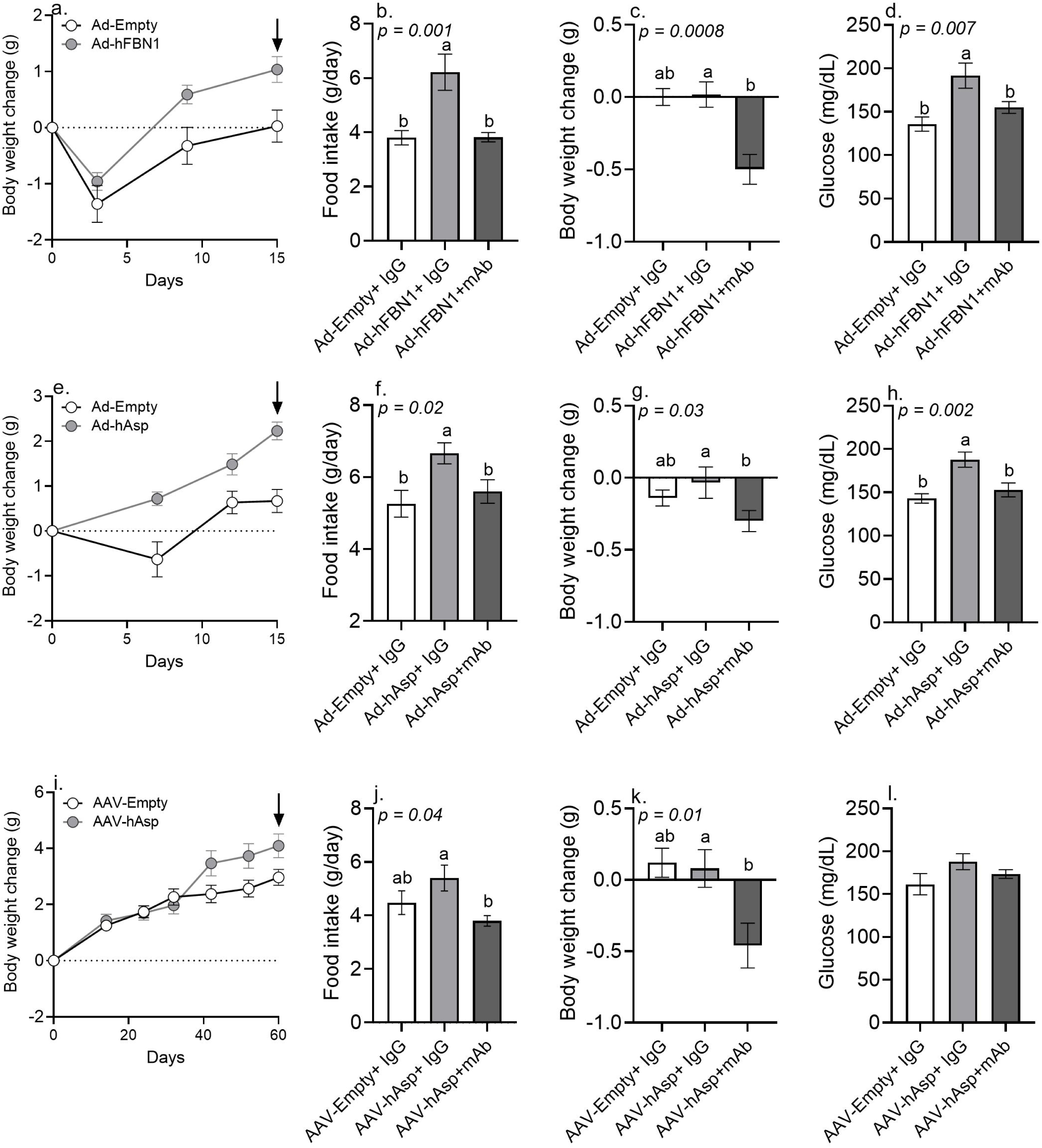
Immunologic neutralization fully rescues the metabolic effects of viral induction of plasma asprosin. (a) Body weight change was measured over 15 days after 12-week-old, male, C57Bl/6 mice were tail-vein-injected with Ad-empty or Ad-FBN1 (3.6 × 10^9^ pfu/mouse, n = 12/group) viruses. Downward arrow indicates the day of mAb treatment described below. (b-d) Cumulative food intake, body weight change, and blood glucose were measured 24 hours after intra-peritoneal injection of indicated control, isotype-matched IgG or anti-asprosin mAbs (n = 6/group) in the above mice. (e) Body weight change was measured over 15 days after 12-week-old, male, C57Bl/6 mice were tail-vein-injected with Ad-empty or Ad-Asprosin (5 × 10^10^ pfu/mouse, n = 12/group) viruses. Downward arrow indicates the day of mAb treatment described below. (f-h) Cumulative food intake, body weight change, and blood glucose were measured 24 hours after intra-peritoneal injection of indicated control, isotype-matched IgG or anti-asprosin mAb (n = 6/group) in in the above mice (i) Body weight change was measured over 60 days after 12-week-old, male, C57Bl/6 mice were tail-vein-injected with AAV8-empty or AAV8-Asprosin (1 × 10^12^ GC/mouse, n = 10/group) viruses. Downward arrow indicates the day of mAb treatment described below. (j-l) Cumulative food intake, body weight change, and blood glucose were measured 24 hours after intra-peritoneal injection of indicated control, isotype-matched IgG or anti-asprosin mAbs (n = 5/group) in in the above mice. Different and same alphabets on bars indicate presence or absence of significant difference, respectively, between groups, as determined by 1-way ANOVA. P < 0.05 considered statistically significant.

When injected with anti-asprosin mAb, mice with viral overexpression of human asprosin restored their food intake to the level observed in empty virus-treated mice (Fig 3b,f,j), and this resulted in an average net body weight loss of 0.3 to 0.5g within the first 24 hours (Fig 3c,g,k). A single injection of mouse anti-asprosin mAb also reduced baseline glucose to levels observed in Ad5-empty injected mice (Fig 3d,h,i). The results of this combined gain-of-function/rescue study indicate that viral overexpression of asprosin is a valuable tool to study asprosin effects in non-obese mice, and that immunologic neutralization fully rescues the metabolic effects of elevated plasma asprosin.

### Chronic asprosin-neutralization mitigates hyperphagia, hyperglycemia and weight gain in obese mice

In DIO mice treated with daily injection for 10 days, we observed a ∼10% decrease in body weight with asprosin neutralization (Fig 4a) that was associated with improved glucose tolerance on day 11 (after 1 day without mAb) (Fig 4b). On day 13 however (10 days of once daily mAb treatment followed by 3 days without mAb treatment), the body weight difference between the two groups remained the same (Fig 4c) but the improved glucose tolerance disappeared (Fig 4d), showing once again that the effect of asprosin neutralization on glucose homeostasis is independent of systemic improvements in metabolism due to weight loss, and also demonstrating the short effect-life of the mAb (24-48 hours).

**Figure 4:**
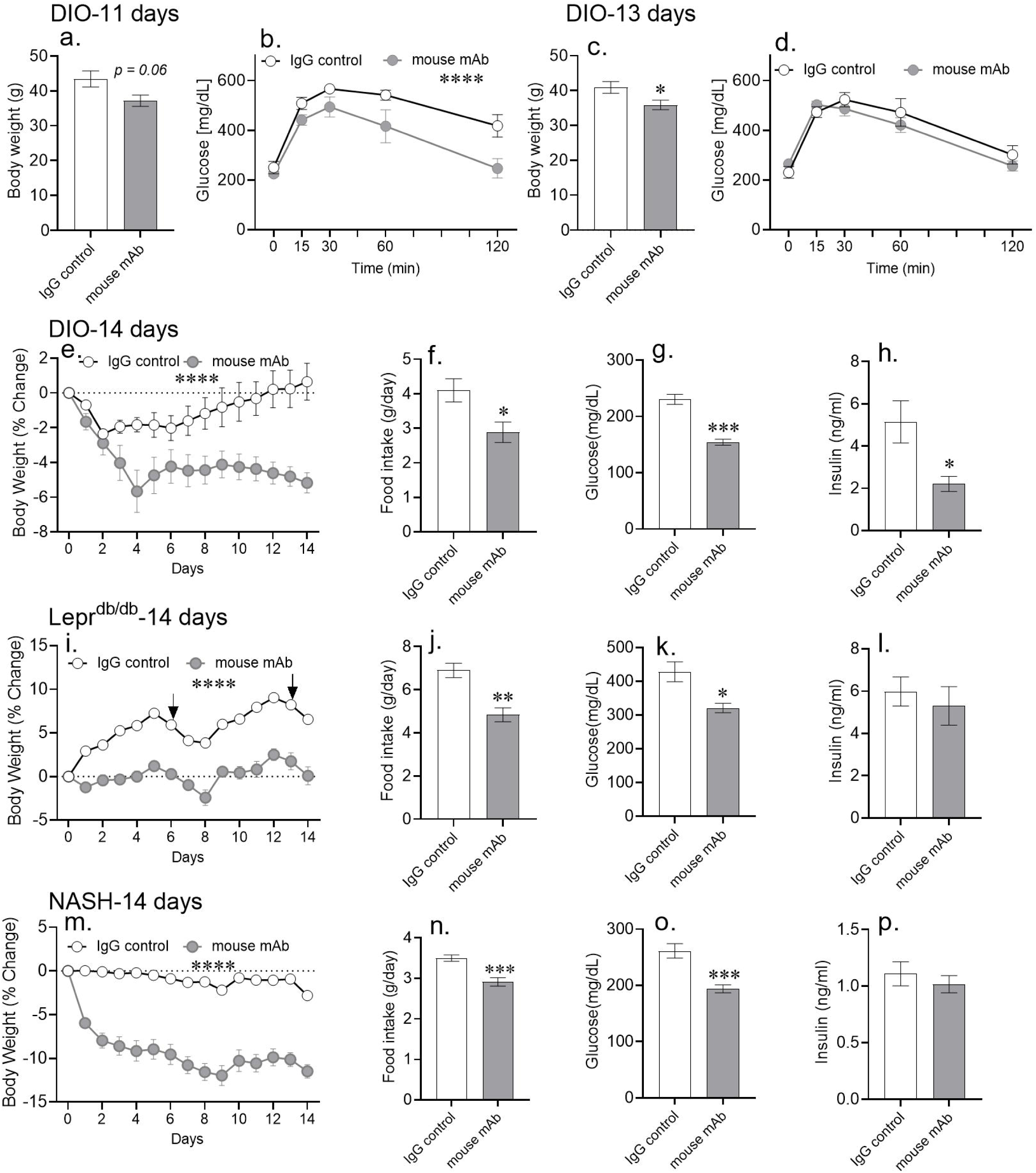
Chronic asprosin-neutralization improves metabolic syndrome in three independent mouse models. (a-d) Body weight and glucose tolerance were measured on day 11 (a,b) and day 13 (c,d) after 10 days of once daily intra-peritoneal injection of control, isotype-matched IgG or anti-asprosin mAb in 16-week-old, male, DIO (diet induced obesity) mice (n = 5/group). (e-p) Percent change in body weight, 24h cumulative food intake (measured on day 7), and blood glucose and plasma insulin (6h post treatment on day 14) levels were measured after 14 days of once daily intra-peritoneal injection of control, isotype-matched IgG or anti-asprosin mAb in 16-week-old, male DIO mice (n = 4 or 5 per group; e-h), 16-week-old, male *Lepr*^db/db^ mice (n = 5 or 6/group; i-l) and 30-week-old, male mice on NASH diet (n = 7 per group; m-p). Asterisk (*) indicate the range of alpha as determined by the t-test (two groups, one time point), or analysis of variance (ANOVA, sets involving multiple groups and time points. * p<0.05, ** p<0.01, *** p<0.001, and **** p<0.0001. Downward arrows in (m) indicate the day of submandibular bleed in *Lepr*^db/db^ mice.

We followed up the single-dose and 10-day studies by testing the effect of asprosin neutralization for 2 weeks. A 14-day course of daily injection of an anti-asprosin mAb significantly reduced food intake and body weight in diet-induced MS (DIO mice; Fig 4e,f), genetic MS (*Lepr*^db/db^ mice; Fig 4i,j) as well as in mice on a NASH diet (Fig 4m,n). On average, mice treated with the mAb weighed approximately 5% - 11% less than mice treated with a control mAb, indicating that the observed reduced food intake results in a net calorie deficit and weight loss. Mice on DIO and NASH diet lost weight, whereas *Lepr*^db/db^ mice were protected from excessive weight gain during the treatment period. Furthermore, while all three mouse models of MS displayed improvement in hyperglycemia (Fig 4g,k,o), a significant reduction in plasma insulin after mAb treatment was observed only in DIO mice (Fig. 4h,l,p).

A single injection of the anti-asprosin mAb significantly reduced plasma levels of total cholesterol, LDL, glycerol and triglycerides in DIO mice, measured 6 hours after injection in mice without access to food (supplementary Fig 4a-f). Chronic treatment for 2 weeks however, in DIO, *Lepr*^db/db^, and NASH diet-treated mice, did not display the uniform improvements in plasma lipids that were noted with a single injection, despite marked improvements in appetite, body weight and plasma glucose levels (supplementary Fig 4g-x). Thus, while the positive effect of chronic asprosin neutralization on appetite, body weight and glucose homeostasis is clear; its positive effect on plasma lipids appears inconsistent. This inconsistency may reflect adaptations and physiological differences in the three mouse models studied.

### A 14-day course of anti-asprosin mAb treatment improves liver health in mice

Inflammation, steatosis, and fibrosis are common hepatic manifestations of MS. We sought to assess the effect of asprosin neutralization on liver health. A significant reduction in mRNA expression of pro-inflammatory markers in anti-asprosin treated DIO mice indicated that hepatic inflammation was markedly suppressed after 14 days of mAb treatment, compared to the control group (Fig 5a). However, prolonged asprosin-neutralization had varied effects on hepatic inflammation in other mouse models of MS. In *Lepr*^db/db^ mice, expression of proinflammatory cytokines, *Il6* and *Tnf*α, was significantly reduced in the anti-asprosin group, but the across-the-board decrease in transcripts of inflammatory candidates observed in DIO mice was not seen in *Lepr*^db/db^ mice, reflecting physiological differences in DIO and *Lepr*^db/db^ models (Fig 5b). In the NASH diet-treated mice, anti-inflammatory *Tgf*β expression was significantly elevated while pro-inflammatory *Ccl2* expression was reduced. Complicating the picture, a significant increase in pro-inflammatory *Fas* and *Cxcl2* gene expression was also noted (Fig. 5c). Thus, in the NASH diet-treated mice the overall effect on tissue inflammation is harder to deduce, and probably reflects differential adaptations to the specialized NASH diet^12,13^.

**Figure 5:**
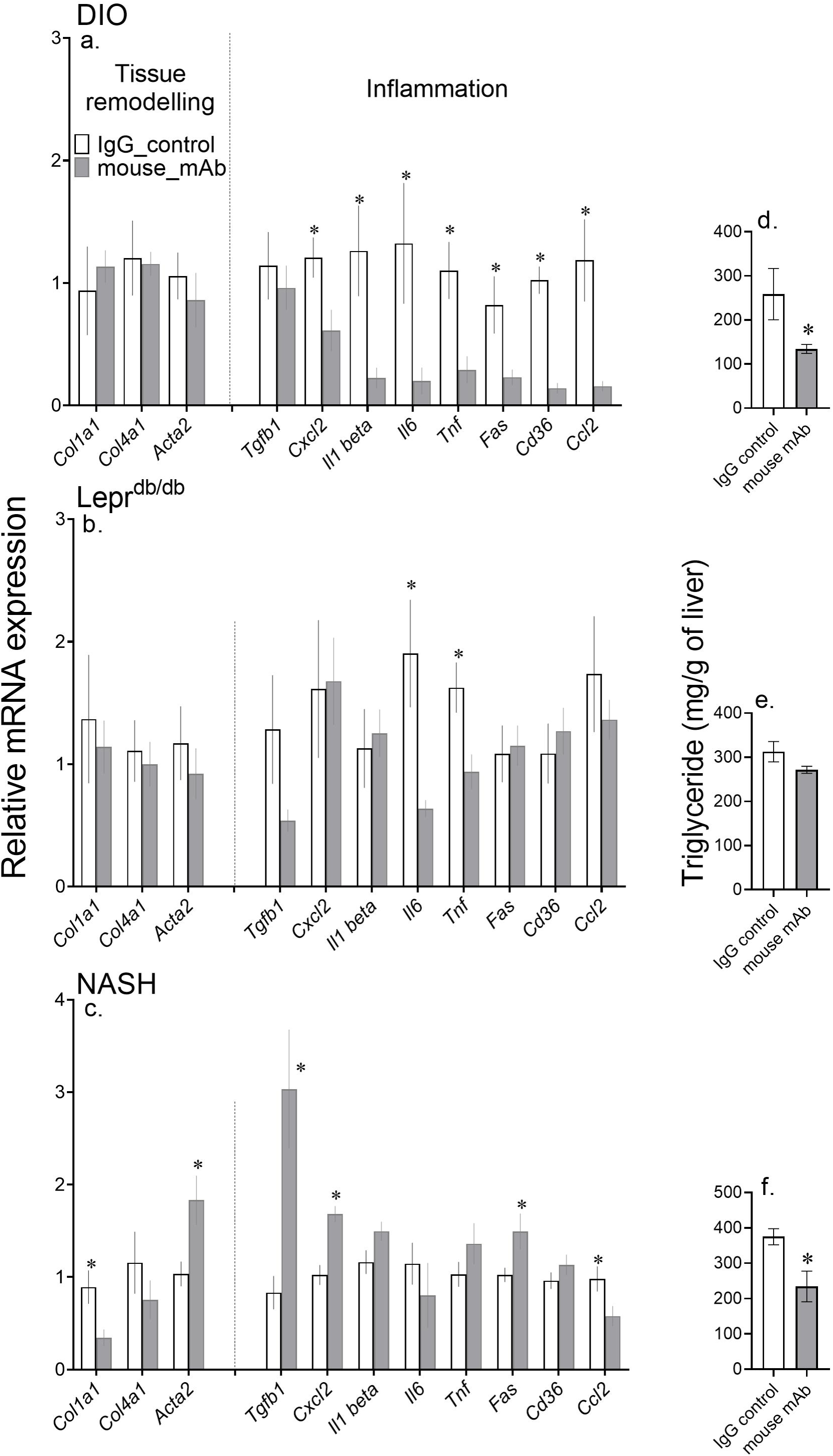
A 14 day course of anti-asprosin mAb treatment improves liver health in MS mouse models. (a-c) mRNA expression levels of genes involved in tissue remodeling (*Col1a1, Col4a1, Acta2*) and inflammatory pathways (*Tgf*β, *Cxcl2, Il1*β, *Il6, Tnf, Fas, Cd36, Ccl2*), and (d-f) triglyceride levels were measured in liver of 16-week-old, male DIO, 16-week-old, male *Lepr*^db/db^ and 30-week-old, male NASH diet fed mice (n = 5-7/group) after 14 days of once daily intra-peritoneal injection of control, isotype-matched IgG or anti-asprosin mAb (250 µg/ mouse). Asterisk (*) indicate the range of alpha; * p<0.05, ** p<0.01, *** p<0.001, and **** p<0.0001, as determined by student T-test.

Interestingly, the expression of genes associated with fibrosis (*Col1a1)* and tissue healing/remodeling (*Acta2)* in NASH diet-treated mice was significantly altered with asprosin neutralization (Fig 5c), suggesting improvement. Hepatic fibrosis and remodeling are only known to occur in mice fed the NASH diet, but not in DIO or *Lepr*^db/db^ mice^12,14-16^. Thus, it is remarkable that asprosin neutralization has an effect on these parameters only in NASH diet-treated mice and not in the other two models of MS. Hepatic triglyceride content was significantly reduced in DIO and NASH diet-treated mice (Fig. 5d,f), indicating reduced lipid burden and improved liver health. This change was not noted in *Lepr*^db/db^ mice (Fig. 5e), likely due to the role of leptin signaling in de novo hepatic lipogenesis^17,18^.

### Neutralization of asprosin using mAbs from distinct species and against distinct epitopes is equally protective

We wondered whether the positive metabolic effects of asprosin neutralization depended on a specific asprosin epitope, or whether multiple epitopes could be targeted. We generated a new rabbit mAb against full-length, glycosylated human asprosin and a new fully human mAb derived from a phage display library. A single injection of each of the three mAbs resulted in significantly improved glucose tolerance (Fig 6a), and a reduction in food intake (Fig. 6b) and body weight (Fig. 6c) in DIO mice. Of these three mAbs, the mouse and human mAbs compete for binding to asprosin and so recognize the same or an overlapping epitope, whereas the rabbit mAb recognizes a distinct epitope, displaying no binding competition (Fig. 6d). This indicates that, insofar as these two epitopes are concerned, the positive metabolic effects of asprosin neutralization are epitope-agnostic. We speculate that formation of an asprosin-mAb complex either accelerates asprosin disposal or prevents ligand-receptor interaction, effectively inhibiting the activity of endogenous asprosin. These results also effectively rule out any potential off-target effects of the mAbs being responsible for their positive effects on metabolic health.

**Figure 6:**
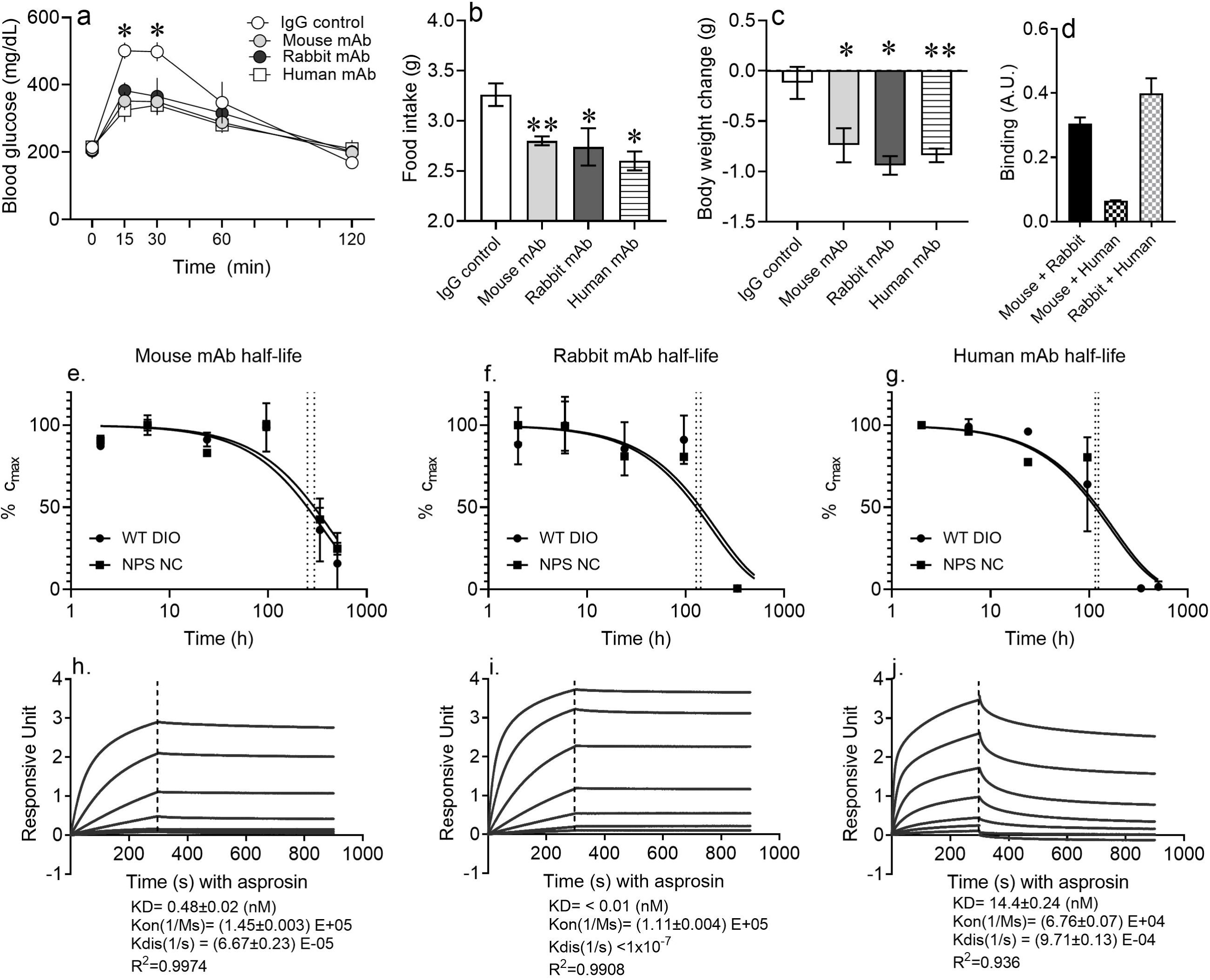
Pharmacokinetics of epitope agnostic anti-asprosin mAbs from different sources. ***(a-c): Neutralization of asprosin using mAbs from different sources is equally protective***. mAb generated using mouse immunization with a 28-mer asprosin peptide (mouse mAb), immunization of rabbits with recombinant full length human asprosin (rabbit mAb) and recombinant mAb generated by panning phages from a naïve human antibody library (human mAb) were injected IP in 16-week-old, male DIO mice (250 µg/mouse, ∼5 mg/kg) and the indicated endpoints measured (n = 5/ group). (d) ***Epitope competition assay***: In a sandwich ELISA, asprosin captured by each mAb (mouse, rabbit or human mAb) was detected by each of the three mAbs in a 3-by-3 matrix to determine competition for their respective epitopes. (e-g) ***Half-life of asprosin-neutralizing mAbs***. Mouse models of ‘high asprosin’ (16-week-old, male mice with diet-induced obesity; DIO), and ‘low asprosin’ (10-week-old, male NPS mice) were injected with mouse, rabbit or human mAb against asprosin 250µg mAb in 500µl 0.9% saline; n = 2/group). mAb levels were determined in mouse plasma collected at 2, 6, 24, 96, 336, and 504h post injection to determine *in vivo* half-life of mAbs. (h-j) ***Anti-asprosin mAb binding affinity***. Equilibrium dissociation constant (KD) of recombinant asprosin binding to anti-asprosin mAb was determined by a 1:1 binding model and use of global fitting method on Pall ForteBio’s Octet RED96 system. Asterisk (*) indicate the range of alpha as determined by analysis of variance (ANOVA). * p<0.05, ** p<0.01, *** p<0.001, and **** p<0.0001.

### Pharmacokinetic parameters of asprosin neutralization

Using an antigen capture ELISA, we measured the plasma concentrations of free mAbs at periodic intervals after injection to calculate their half-life in both endogenously high-asprosin (DIO mice) and low-asprosin (NPS mice) conditions (Fig. 6e-g). No mAbs against asprosin were detected before injection, whereas a calculated peak concentration of 0.1 mg/ml ligand-free mAb was observed two hours after injection. In DIO mice, the free mouse anti-asprosin mAb had the longest half-life, of approximately 11 days, the free rabbit anti-asprosin mAb had a half-life of approximately 5.6 days, and the free human anti-asprosin mAb had a half-life of approximately 3.2 days. Of note, the half-lives of these three mAbs were fairly similar in NPS mice, indicating that the concentration of circulating asprosin does not appear to directly influence the rate of mAb clearance.

Asprosin-mAb affinity was measured on the Octet RED96 system (Fig. 6h-j). The K_D_ of the rabbit mAb to recombinant human asprosin was calculated as < 10 pM from a k_on_ of 1.11 × 10^5^ M^−1^s^−1^ and a k_off_ <1×10^−7^ s^−1^ (r^2^ = 0.99, χ^2^ = 99.88). The mouse mAb displayed a K_D_ of 0.48±0.02 nM with k_on_ of (1.45±0.003) × 10^5^ M^−1^s^−1^ and k_off_ of (6.67±0.23) × 10^−5^ s^−1^ (r^2^ = 0.99, χ^2^ = 15.05). The human mAb displayed a K_D_ of 14.4±0.24 nM with k_on_ of (6.76±0.07) × 10^4^ M^−1^s^−1^ and k_off_ of (9.71±0.13) × 10^−4^ s^−1^(r^2^ = 0.94, χ^2^ = 408.8). Thus the rabbit mAb has the highest affinity for asprosin and the human mAb the lowest, over a 1000-fold range. Nevertheless, all three mAbs were able to functionally neutralize asprosin *in vivo*.

## Discussion

We recently discovered a novel glucogenic and orexigenic hormone, named asprosin, whose circulating levels are elevated in mice, rats, and humans with metabolic syndrome^2-8^.

In this report, we show that three independent asprosin gain-of-function tools (Ad5-*FBN1*, Ad5-*Asprosin*, AAV8-*Asprosin*) produce an increase in appetite, body weight and blood glucose, and three independent asprosin loss-of-function tools (mouse, rabbit and human mAbs against at least two distinct asprosin epitopes) produce the opposite *in vivo*, compared with their respective controls. We were even able to demonstrate mAb-induced rescue of the viral-induced plasma asprosin elevation, lending a very high level of solidity to our conclusions. Additionally, two independent groups have now recapitulated our original study on the discovery of asprosin in the process of elucidating the hepatic asprosin receptor^3^ and an asprosin-like hormone^19^. This is particularly important given the inability of one group to recapitulate our original study^20^, most likely due to use of poor quality recombinant asprosin. Recombinant asprosin remains an unreliable research tool, even in our hands, and will likely remain so until the biochemical factors (potential chaperons for example) that govern asprosin stability and activity come to light. The viral vectors and mAbs reported in this study will be made freely available to the research community to study asprosin biology *in vivo*.

Asprosin neutralization resulted in reduced food intake and body weight in mice on a high fat diet (diet induced obese, DIO mice) as well as in more severe preclinical models of MS, such as mice on a NASH-inducing diet and mice with a genetic loss of leptin signaling (*Lepr*^db/db^). These results indicate that asprosin neutralization is effective independent of leptin, and opens therapeutic avenues for a wide range of cases of hyperphagia and obesity. Concurrent with decrease in appetite and body weight, anti-asprosin mAb therapy improved the MS-associated hyperglycemia in all preclinical models studied. Importantly, a low IC_50_ of 30-55 µg/mouse (∼1.5mg/kg) was determined for the four key features of MS (hyperphagia, obesity, hyperglycemia and hypertriglyceridemia). The low IC_50_, well within the spectrum of acceptable therapeutic mAb dosing, highlights the pharmacological inhibitory potency of asprosin neutralization for MS in its entirety, rather than improving only a particular manifestation of it^21^. While the effect of escalating dose on appetite reduction is linear, it exhibits a defined upper and lower threshold when it comes to improvement of other attributes of MS such as blood glucose, insulin, triglycerides and body weight. This may indicate arrival at a physiological equilibrium state and the impact of compensatory mechanisms to prevent further, potentially disastrous changes.

Interestingly, mAb treatment did not result in any changes in lean euglycemic WT mice with the exception of a short-lived reduction in blood glucose (without crossing the threshold for overt hypoglycemia). This result suggests the dependence of anti-asprosin mAb therapeutics on high endogenous asprosin levels, and portends a strong safety profile.

A single dose of the anti-asprosin mAb improved MS-associated dyslipidemia in DIO mice as evidenced through a reduction in total cholesterol, LDL, triglycerides and glycerol. This could potentially be explained by the glucose and insulin lowering effects of asprosin neutralization leading to suppression of lipogenesis^22,23^. With chronic neutralization reductions in total cholesterol were noted in DIO and *Lepr*^db/db^ mice. However, the effect on other lipid species was not as uniform as with single dosing, suggesting compensatory adaptations with chronic asprosin loss-of-function.

DIO mice on an anti-asprosin therapy for 2 weeks displayed remarkable improvements in hepatic inflammation, as evidenced by reductions in chemokine and cytokine expression, and steatosis. In mice on a NASH diet however, the effects of asprosin neutralization on hepatic inflammation were mixed. A significant reduction in *Ccl-2* expression indicated that infiltration of myeloid cells into the liver was abrogated^24^. However, there was an increase in pro-inflammatory *Fas* and *Cxcl2* expression, and an increase in the anti-inflammatory *Tgf-*β. Thus, the overall effect on inflammation in NASH mice is unclear. This may reflect the intensity of inflammation produced by the NASH diet, which is greater than that produced by a simple high fat diet^12,13^. Advanced stages of NASH, in particular fibrosing NASH, are a leading cause of liver cirrhosis in MS patients^12^. In response to asprosin neutralization in mice on a NASH diet, we noted decreased expression of genes associated with hepatic fibrosis such as *Col1a1*. We also noted an increased expression of *Acta2*, a marker of hepatic wound healing via stellate cell to liver-specific myofibroblast transformation and increased cellular motility^15,16^. Taken together, these results suggest that anti-asprosin mAbs are protective against, and could potentially reverse, MS associated liver damage.

In addition to the mouse mAb we generated rabbit and fully human mAbs for this study. The human mAb shares an epitope with the mouse mAb, while the rabbit mAb recognizes a distinct asprosin epitope. All three mAbs at a dose of 250 µg/mouse (∼5 mg/kg) improved glucose tolerance and resulted in reduced food intake and body weight. This suggests that steric inhibition of asprosin’s interaction with its receptor is either epitope-agnostic, or that the beneficial effects of asprosin neutralization depend on rapid asprosin disposal, or some combination of these two possibilities. Further, all three mAbs showed an equilibrium dissociation constant (K_D_) in the picomolar to low nanomolar range indicating high affinity of mAb-asprosin binding^25^. Despite the high affinity and long half-life of the mouse mAb, maintenance of its effects required daily administration. A short “effect-life,” out of proportion to mAb half-life, has previously been reported for other mAbs against various circulating antigens^26-28^. This short effect-life might be explained by high asprosin-mAb complex stability under various physiological conditions, leaving newly produced asprosin uninhibited^28,29^, thereby necessitating new mAb administration for continued pharmacological effect. In other words, there is sufficient precedence for this issue when it comes to circulating antigens and overcoming this requires a variety of mAb engineering approaches, which are under consideration for anti-asprosin mAbs at this time.

Conceptually however, the studies presented here offer promise for targeting asprosin in the treatment of MS, with planned lead optimization (to improve effect-life and humanize the lead candidate), exploratory toxicity and efficacy studies in other species such as nonhuman primates, potentially leading to human trials. In addition, although the receptor for asprosin in the liver has been discovered recently^3^, the identity of its receptor in the CNS remains unknown. These receptor discovery efforts are essential for the development of orally-bioavailable, small molecule inhibitors of the asprosin pathway, which are validated by the ligand-targeting mAb approach presented here.

In summary, we provide evidence that demonstrates the potential of asprosin neutralization using multiple mAbs *in vivo* to treat MS. As opposed to current and past therapies that target satiety, this approach directly inhibits appetite and separately reduces the blood glucose burden by inhibiting hepatic glucose release. Thus, it is a dual-effect therapy that targets the two pillars of MS, over-nutrition and increased glucose burden. The results presented here, pre-clinically validating the pharmacological inhibition of asprosin for the treatment of MS, are a source of high optimism moving forward.

## Methods

### Mice

12 week old C57Bl/6J (wild-type, WT) mice, leptin-receptor deficient mice (*Lepr*^db/db^), and 16-week-old mice with diet-induced obesity (DIO) were purchased from Jackson Laboratories. Mice were fed normal chow (5V5R, Lab Supply), dustless pellet diet (F0173, Bio-Serv), 12 weeks of high fat diet (60% calories from fat, TD.06414, Envigo Teklad), or 24 weeks of NASH-high fat diet (AMLN-diet, D09100301, Research Diets), where indicated. Prior to all interventions, mice were standardized across groups ensuring equal distribution of body weight for the various treatment groups.

### mAb generation

A majority of the studies were conducted with a mouse mAb (M1). M1 was generated using traditional hybridoma techniques by immunizing mice with a 28 amino acid peptide KKKELNQLEDRYDKDYLSGELGDNLKMK located close to the C-terminus of asprosin. Where indicated only, rabbit and fully-human anti-asprosin mAbs were used in parallel with the mouse anti-asprosin mAb. The rabbit mAb was generated by immunizing rabbits with recombinant full-length human asprosin at RevMAb Biosciences, USA, and cloning variable region genes from positive single memory B cells based on protocols described previously.^30^ A fully human mAb was generated from a naïve human phage display antibody library by panning against recombinant full-length human asprosin (Texas Therapeutics Institute at the University of Texas Health Science Center at Houston). All three mAbs display cross-reactivity to mouse and human asprosin.

### mAb injection

Mice received either a single dose of 250µg/mouse (∼5-6 mg/kg) mAb intraperitoneally in 500µl USP grade saline, or repeated daily dose for up to 14 days. Injections were performed between 9am and 11 am both for single and repeated dose studies. For dose response curve, DIO mice received a single IP injection of different doses of control, isotype-matched IgG or anti-asprosin mouse mAb (15.63, 31.25, 62.5, 125, and 250µg/mouse, corresponding to 0.4, 0.85, 1.64, 3.28 and 6.88 mg/kg; n = 5/dose) at 11 to 11:30 am.

### Adenovirus and Adeno-associate virus experiments

12-week-old C57Bl/6J mice were injected intravenously via the tail-vein with adenovirus (Ad5) or adeno-associated virus, serotype 8 (AAV8) containing the human FBN1 or his-tagged human asprosin coding region. Mice injected with Ad5-empty and AAV8-Empty served as controls for experimental mice (detailed procedure in Suppl. Methods).

### mRNA expression

Whole livers were isolated and flash frozen in liquid nitrogen. A small aliquot was processed for total RNA extraction using RNeasy Mini spin columns (Qiagen). cDNA was synthesized using a high capacity RNA-to-cDNA kit (Thermo Fisher Scientific) and gene expression was measured using gene specific primers (Supplementary Methods) and a probe-based TaqMan assay (Thermo Fisher Scientific). Expression data was calculated using the ΔΔCt method and normalized to a control sample.

### Half-life of asprosin-neutralizing mAbs

For assessing *in vivo* half-life of mAbs, mouse models of ‘high asprosin,’ C57Bl/6J (wild-type, WT) mice with diet-induced obesity (DIO, n = 6), and ‘low asprosin’ (NPS mice, n = 6) were injected with mouse mAb (n = 2/group), rabbit mAb (n = 2/group), or human mAb (n = 2/group) (250µg mAb in 500µl 0.9% saline). Thereafter, mAb levels were determined in mouse plasma collected at 2, 6, 24, 96, 336, and 504h post injection (detailed procedure in Suppl. Methods).

### Epitope Competition

To determine whether the 3 mAbs recognized overlapping or distinct epitopes, we captured recombinant asprosin (20 nM) on an ELISA plate coated with one of the three mAbs (100ng/well), followed by detection with each of the three mAbs in a 3 × 3 matrix format (100ng/well, detailed in Suppl. Methods).

#### Asprosin-mAb binding affinity measurement with BLI

Asprosin-mAb affinity was measured on Pall ForteBio’s Octet RED96 system using Ni-NTA biosensor (ForteBio, Cat#18-5102). K_D_ was determined from 7 kinetic curves (detailed procedure in Supplementary Methods) fitted by a 1:1 binding model and use of global fitting method in ForteBio’s data analysis software.

### Statistical analysis

Data was graphed and analyzed using GraphPad Prism (Version 6 and higher). Data is presented as mean +/-standard error of the mean. Depending on the format of data, analysis was by using t-test (two groups, one time point), or analysis of variance (ANOVA) sets involving multiple groups and time points. ANOVA analysis is indicated with the word “ANOVA” in figures next to the significance asterisk). Non-linear four parametric variable slope (least square regression) analyses determined the IC_50_ and R^2^ values in dose response studies. Significance is presented using the asterisk symbol (* p<0.05, ** p<0.01, *** p<0.001, and **** p<0.0001). In situations where a non-significant trend was observed, the full p-value is presented.

## Acknowledgments

We thank Georgina Salazar, Andrew Pieper, Richard Premont, Mukesh Jain and Jonathan Stamler for critical reading of the manuscript. This work was supported by the Cancer Prevention and Research Institute of Texas (RP150551 and RP190561), the Welch Foundation (AU-0042-20030616 and I-1834), the NIDDK (DK102529, DK118290) and the Harrington Discovery Institute.

## Author Contributions

A.R.C. conceptualized the study; I.M., C.D., E.S.S, Z.K., W.X., J.H., W.X. and N.Z. performed the experiments; A.R.C. and Z.A. provided resources; and I.M. and A.R.C. wrote the manuscript.

## Figure legends

**Supplementary Figure 1: *Higher blood glucose and insulin levels in metabolic syndrome patients are associated with elevated asprosin levels***. Blood glucose, plasma insulin and asprosin levels were measured in metabolic syndrome male patients (n = 10; BMI > 25) and age-matched male subjects with normal BMI (< 25). Asterisk (*) indicate the range of alpha; * p<0.05, ** p<0.01, *** p<0.001, and **** p<0.0001, as determined by student T-test.

**Supplementary Figure 2: *Acute asprosin-neutralization reduces blood glucose, but not appetite and body weight in lean mice***. Cumulative food intake and body weight change (at 24h post treatment), baseline blood glucose (at hour 2, 4 and 6 post mAb-treatment) and plasma insulin (at 6h post treatment) was measured after a single dose of anti-asprosin mAb (250µg/mouse) in 12-week-old, male, C57BL/6J lean mice, n = 5 per group. Asterisk (*) indicate the range of alpha; * p<0.05, ** p<0.01, *** p<0.001, and **** p<0.0001, as determined by student T-test.

**Supplementary Figure 3: *Viral overexpression of human asprosin results in a MS-like phenotype in lean mice***.

(a-d) Body weight change, cumulative food intake, blood glucose, and plasma insulin were measured 15 days after 12-week-old, male, C57Bl/6 mice were tail-vein-injected with Ad-empty or Ad-FBN1 (3.6 × 10^9^ pfu/mouse, n = 5/group) viruses.

(f-i) Body weight change, cumulative food intake, blood glucose, and plasma insulin were measured 15 days after 12-week-old, male, C57Bl/6 mice were tail-vein-injected with Ad-empty or Ad-Asprosin (5 × 10^10^ pfu/mouse, n = 12/group) viruses.

(k-n) Body weight change, cumulative food intake, blood glucose, and plasma insulin were measured 57 days after 12-week-old, male, C57Bl/6 mice were tail-vein-injected with AAV8-empty or AAV8-Asprosin (1 × 10^12^ GC/mouse, n = 10/group) viruses.

(e,f,o) Human asprosin levels detected in plasma of Ad-FBN1, Ad-Asprosin and AAV8-Asprosin injected mice is plotted relative to the average background signal detected in Ad-empty and AAV8-empty injected mice.

Asterisk (*) indicate the range of alpha; * p<0.05, ** p<0.01, *** p<0.001, and **** p<0.0001, as determined by student T-test.

**Supplementary Figure 4: *Asprosin-neutralization improves dyslipidemia in MS mouse models***.

(a-f) Plasma levels of total cholesterol (TC), low density lipoproteins (LDL), high density lipoprotein (HDL), triglycerides (TG), free fatty acids (FFA) and glycerol were measured 6h after intra-peritoneal injection of indicated control, isotype-matched IgG or anti-asprosin mAb in 16-week-old, male DIO mice (250µg/mouse, n = 5/group; a-f),

(g-x) Plasma levels of TC, LDL, HDL, TG, FFA and glycerol were measured 24h after 14 days of once daily intra-peritoneal injection of indicated control, isotype-matched IgG or anti-asprosin mAb (250µg/mouse) in 16-week-old, male DIO mice (n = 6/ group; g-l), 16-week-old, male *Lepr*^db/db^ mice (n = 7/group; Db/Db; m-r) and 30-week-old, male mice on NASH diet (n = 7/group; s-x).

Asterisk (*) indicate the range of alpha; * p<0.05, ** p<0.01, *** p<0.001, and **** p<0.0001, as determined by student T-test.

