## Supplementary material for "Asprosin Neutralizing Antibodies as a Treatment for Metabolic Syndrome": Suppl. Methods

### Supplementary Methods

### *Food intake*

Mice were placed in CLAMS metabolic cages (Columbus Instruments) and acclimated for 3 days. Food intake was recorded for 24 hours and mice were returned to normal housing. Food intake was manually measured in 24h experiments in DIO mice and Ad5 and AAV8 studies. In Ad5 and AAV8 studies, mice were singly housed in standard caging and fed a pelleted, dustless diet (F0173, Bio-Serv). In single day DIO experiments, mice were acclimated to crushed high fat diet (60% calories from fat, TD.06414, Envigo Teklad) in single housing. The diet was replenished, weighed, and re-weighed after 24 hours to establish food intake.

### *Adenovirus and Adeno-associate virus experiments*

12-week-old C57Bl/6J mice were injected intravenously via the tail-vein with adenovirus (Ad5) or adeno-associated virus, serotype 8 (AAV8) dissolved in 150 µl USP grade sterile saline. Mice injected with Ad5-empty (3.6 x 10^9^ pfu/ mouse) served as controls for experimental mice that received Ad5-FBN1 virus (3.6 x 10^9^ pfu/ mouse) containing the human FBN1 coding region under control of a CMV promoter. Mice injected with Ad5-empty (5 x 10^10^ pfu/mouse) served as controls for experimental mice that received Ad5-IL2-Asprosin (5 x 10^10^ pfu/mouse) containing an N-terminal his-tagged human asprosin coding region preceded by an IL2 signal peptide, under control of an EF1 promoter. Mice injected with AAV8-Empty (1 x 10^12^ GC /mouse) served as controls for experimental mice that received AAV8-IL2-Asprosin (1 x 10^12^ GC /mouse) containing an N-terminal his-tagged human asprosin coding region preceded by an IL2 signal peptide, under control of an EF1 promoter.

### *Plasma metabolic parameters*

Mouse glucose was determined using a hand-held glucometer (OneTouch Ultra2, LifeScan) from a droplet of tail blood and insulin was measured using Crystal Chem (catalog # 90080) mouse insulin ELISA kit. Mouse plasma total cholesterol, HDL (high-density lipoprotein cholesterol), LDL (low-density lipoprotein cholesterol), triglyceride (TG), free fatty acids (FFA), glycerol and liver triglyceride levels were measured by the mouse metabolism and phenotyping core at Baylor College of Medicine, Houston, Texas. Glucose and insulin levels in human plasma purchased from BioIVT were measured using a chromogenic glucose quantitation kit (Cayman Chemicals; catalog # 10009582) and a human insulin ELISA kit (Raybiotech; catalog # ELH-Insulin), respectively.

***Primer sequences:***

*Col1a*- F: ccgctggtcaagatggtc; R: ctccagcctttccaggttct; *Col4a1*- F: ttaaaggactccagggaccac, R : cccactgagcctgtcacac*, Acta2:* F: *ctctcttccagccatctttcat, R: tataggtggtttcgtggatgc; Tgfβ*- F: tggagcaacatgtggaactc, R: gtcagcagccggttacca; *Cxcl2*- F: aaaatcatccaaaagatactgaacaa, R: ctttggttcttccgttgagg; *Il1β*: ttgacggaccccaaaagat, R: gaagctggatgctctcatctg; *Il6*- F: gctaccaaactggatataatcagga, R: ccaggtagctatggtactccagaa; *Tnf*- F: ctgtagcccacgtcgtagc, R: ttgagatccatgccgttg; *Fas*- F: gcagacatgctgtggatctg, R: tcggagatgctattagtaccttgag; *Cd36*- F: ttgtacctatactgtggctaaatgaga, R: cttgtgttttgaacatttctgctt; *Ccl2*- F: catccacgtgttggctca, R: gatcatcttgctggtgaatgagt.

***ELISA procedures:***

For assessing *in vivo* half-life of antibodies, antibody in plasma was captured on ELISA plate coated with asprosin (100ng/well) and levels of mouse mAb, rabbit mAb and human mAb were detected with HRP conjugated anti-mouse (1:10,000), anti-rabbit (1:10,000) or anti-human 1:10,000) secondary antibodies, respectively.

For detection of endogenous asprosin in human plasma samples and adeno- and adeno-associated virus generated human asprosin in mouse plasma, a human asprosin sandwich ELISA was custom built using mouse monoclonal anti-asprosin antibody against human asprosin amino acids 106–134 (human proﬁbrillin amino acids 2838–2865) as the capture antibody, and a rabbit anti-asprosin monoclonal antibody as the detection antibody. An anti-rabbit secondary antibody linked to HRP was used to generate a signal, and mammalian-cell produced recombinant human asprosin was used to generate a standard curve. Human asprosin was detected in 10 ul of EDTA treated human plasma. For detection of human asprosin in plasma of mice treated with human asprosin expressing adeno- and adeno-associated viruses, plasma samples were first processed for IgG and albumin removal using Proteome purify2 columns (R & D systems, Catalog # IDR002) and concentrated using vivaspin 500 (VS0131) PES filters before running the ELISA.

Epitope competition assay was done to determine whether the 3 mAbs recognized the same epitope or different epitopes. For this, we captured recombinant asprosin (20 nM) on an ELISA plate coated with one of the three mAbs (100ng/well), followed by a detection with each of the three mAbs (100ng/well, 3x3 matrix, detailed methods in Supplementary File), mouse, rabbit or human mAb. Subsequently, captured asprosin was then incubated in parallel with each of the three different mAbs (100ng/well, 3x3 matrix). Depending on which species antibody was used in the second step, a species-matched secondary detection antibody was used. For example, each well that was incubated with a mouse anti-asprosin mAb in the second step was then incubated with an anti-mouse secondary HRP-conjugated antibody. Mouse/mouse, rabbit/rabbit, and human/human pairs served as internal positive control. For clarity, only one version of each combination is presented here, but the competition worked both ways.

***Asprosin-antibody binding affinity measurement with BLI*:**

Antibody affinity was measured on Pall ForteBio’s Octet RED96 system. Recombinant human asprosin (25 mg/mL) was loaded onto a Ni-NTA biosensor (ForteBio, Cat#18-5102) for 300 seconds. Following 20 seconds of baseline in kinetics buffer (ForteBio Cat# 18-5032), the loaded biosensor was dipped in parallel into a series of antibody solutions on increasing concentrations (0.14–300 nM) for 300 seconds to record the association kinetics and then dipped into the kinetic buffer (ForteBio Cat# 18-5032) for 600 seconds to record the dissociation kinetics. Kinetic buffer without antibody was set to correct the background. K_D_ was determined from 7 kinetic curves fitted in a 1:1 binding model using global fitting in ForteBio’s data analysis software.
