## Supplementary figures and images for "Asprosin Neutralizing Antibodies as a Treatment for Metabolic Syndrome"

### Suppl. Figure 1

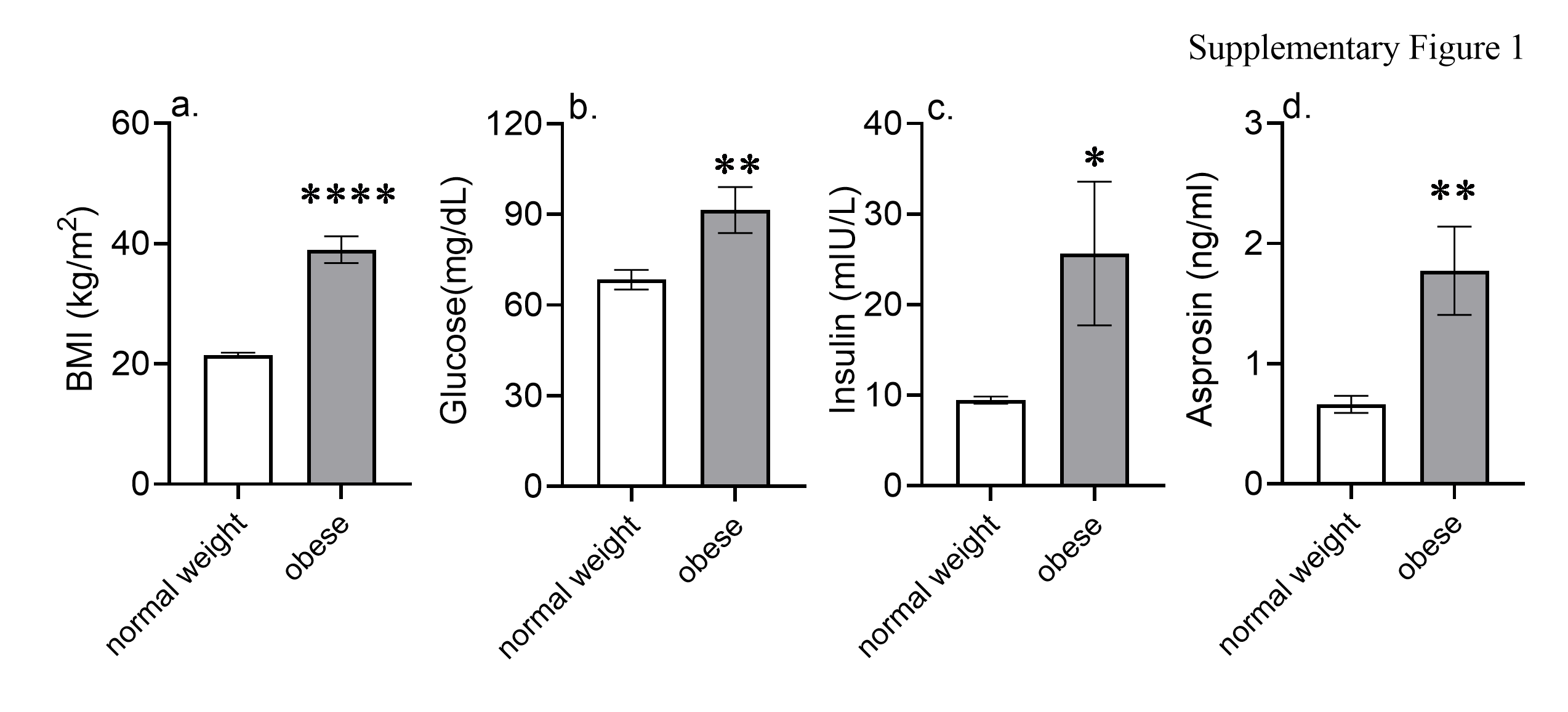

### Suppl. Figure 2

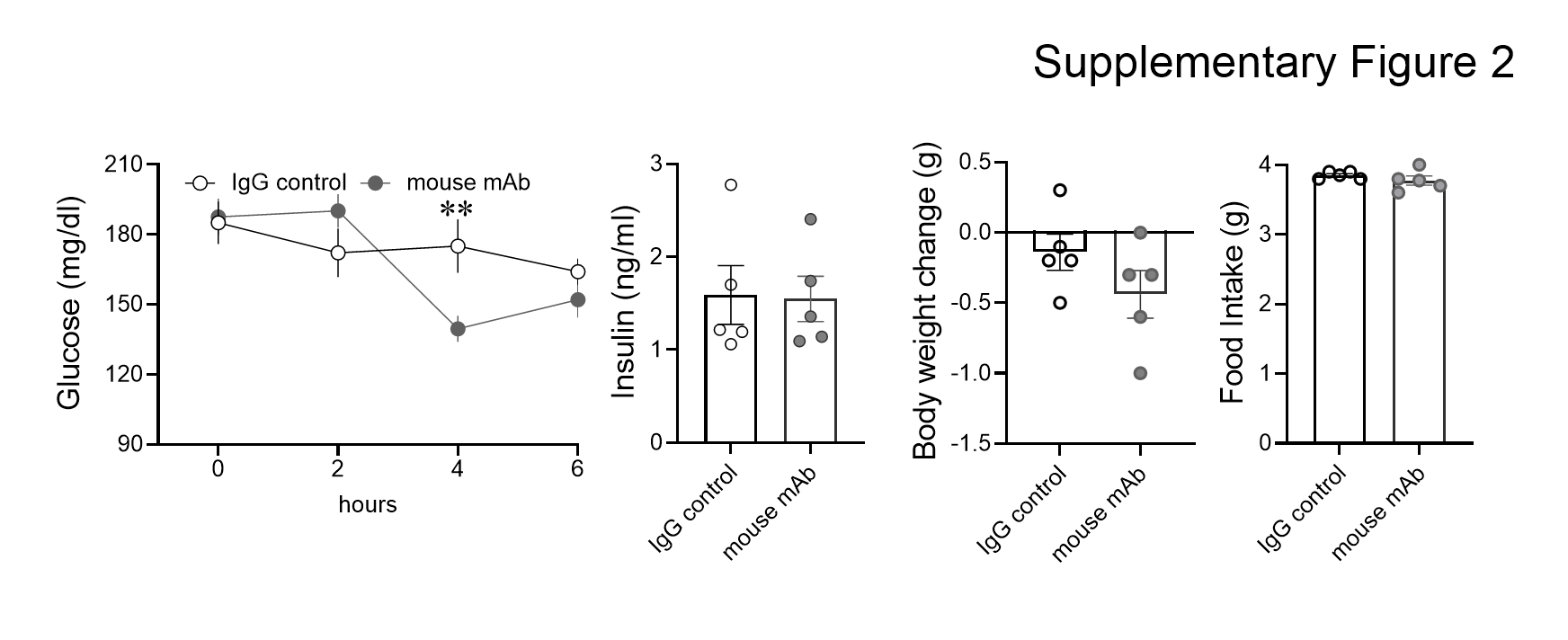

### Suppl. Figure 3

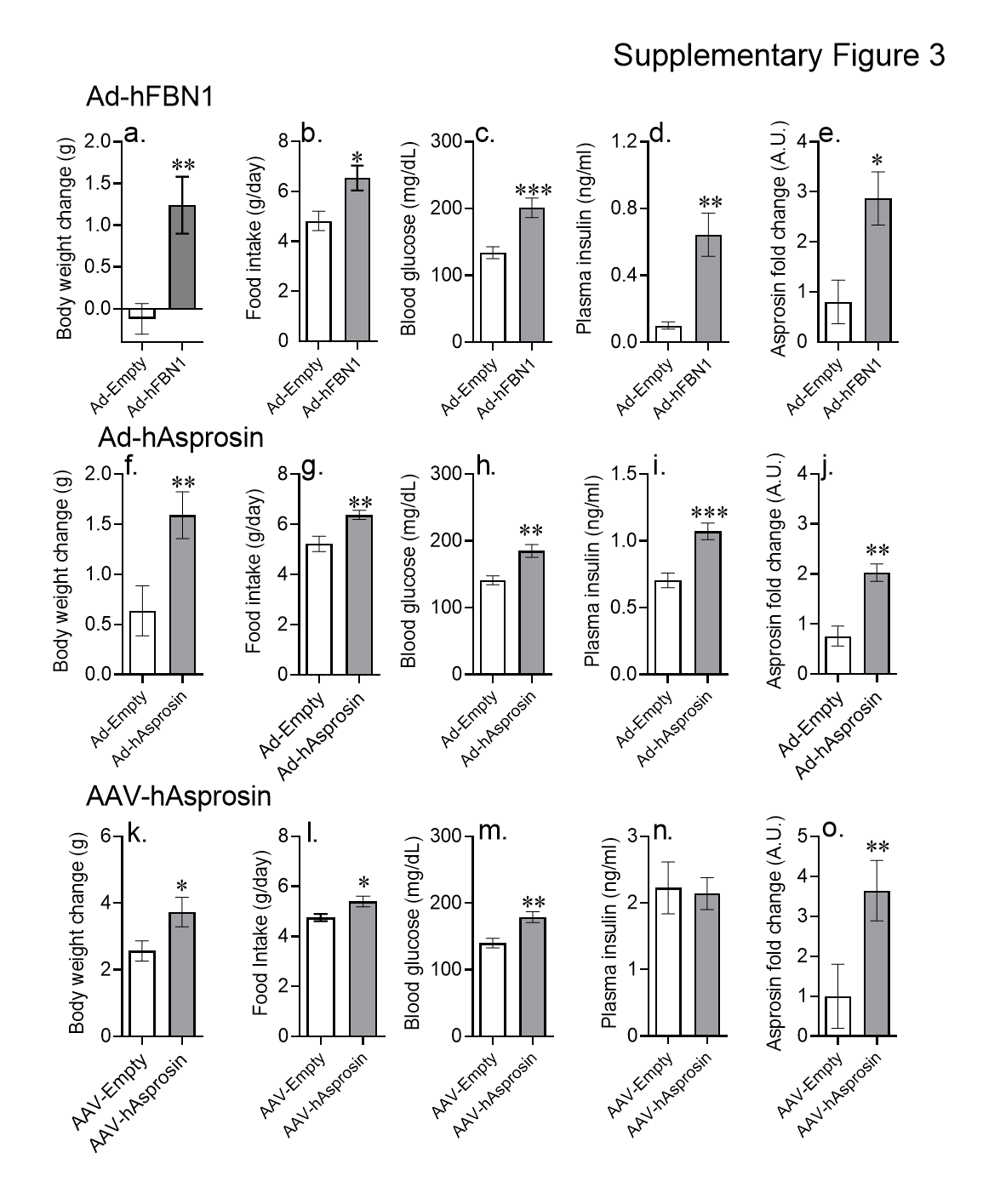

### Suppl. Figure 4

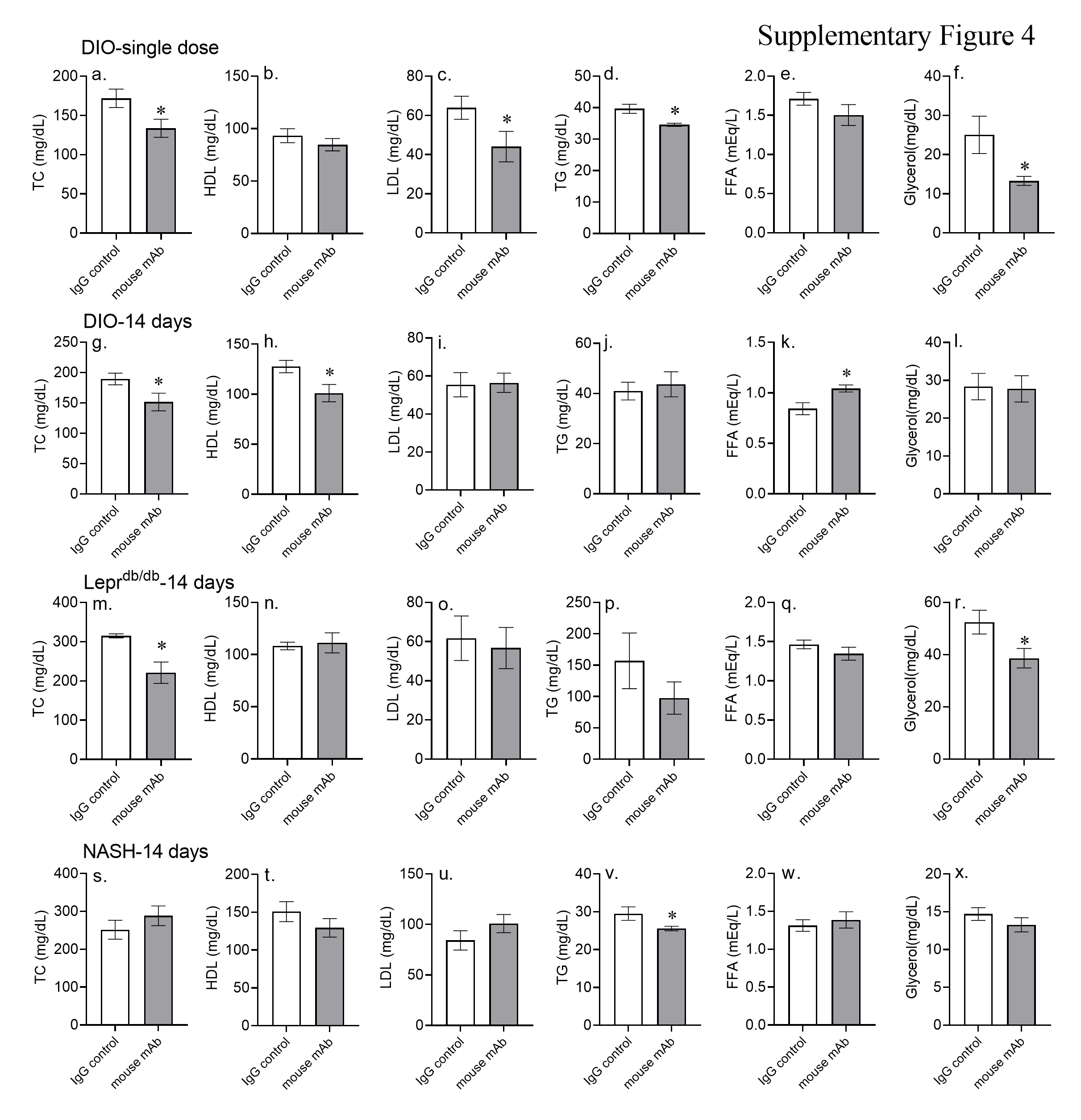
